# High content analysis methods enable high throughput nematode discovery screening for viability and movement behavior in a multiplex sample in response to natural product treatment

**DOI:** 10.1101/432195

**Authors:** Jennifer M. Petitte, Mary H. Lewis, Tucker K. Witsil, Xiang Huang, John W. Rice

**Author notes:** All authors had substantial roles in conceptualization, method development, data acquisition, experimental execution, and data annotation, data analysis and curation. JM Petitte in particular was instrumental in method development, worm husbandry, and scripting of worm analysis algorithms. MH Lewis developed the methodology, sample tracking and data curation to generate the microbial sample test library. TK Witsil wrote all of the necessary python scripts to automate the final data annotation. X Huang was responsible for group supervision, project administration, and development of nematode husbandry methods. JW Rice was responsible for high content method development, results validation and visualization, and original draft preparation. All authors are employees of Novozymes North America, Inc., 108 TW Alexander Drive Bldg. 1A, Durham, NC 27519.

## Abstract

Monitoring nematode parasite movement and mortality in response to various treatment samples usually involves tedious manual microscopic analysis. High Content Analysis instrumentation enables rapid and high throughput collecting of large numbers of treatment data on huge numbers of individual worms. These large sample sizes and increased sample diversity result in robust, reliable results with increased statistical significance. These methods would be applicable to relevant human, crop, or animal worm parasites.

## Introduction

Nematode infection of commercial crops resulting in significant yield loss is a wide-spread economic and food security issue. Control of these crop pathogens has been hampered by development of resistance to pesticides by the targeted pest nematode populations and removal from the market of chemical pesticides due to general environmental toxicity^1^. For these reasons, much of the research targeting methods of regulation of nematodes has shifted focus to alternative control strategies that utilize natural soil microbial and fungal strains^2,3^.

Phenotypic nematode screening methods for active small molecules or microbes capable of controlling nematode infestations in both plants and animals can be as simple as manual microscopic examination of individual samples or as complex as movement-based analysis of high definition video captured worm images and microfluidics approaches^4–7^. All these approaches utilize visualization of some form, either manual scoring of non-movement vs. movement in single samples, or exact single worm movement measurements based on image analysis, such as “The Worminator” system^8^. But typically, plant nematode screens are low-throughput, labor intensive and “mortality” is measured as lack of movement, but not distinguishable from temporary paralysis or true mortality. By combining the sample throughput and ability to capture time resolved images with the accuracy of measuring viable dye influx and concentration on an individual worm basis, we have designed a phenotyping assay method that utilizes the statistical power of high content imaging and the ability to test large sample libraries to screen a proprietary microbial sample library to determine which individual microbe may display possible positive control effects. This effort targeted two widespread crop pests, the Root Knot Nematode *Meloidogyne incognita* (RKN) and the Soybean Cyst Nematodes *Heterodera glycines* (SCN).

## Materials and Methods

### Materials

Original Nematode samples were obtained from the laboratory of Dr. Rick Davis, located at North Carolina State University. Williams 82 soybeans were from the laboratory of Dr. Mary Ann Quade, located at the University of Missouri, and cucumber seeds were Marketmore 76 obtained from JonnySeeds.com. PKH26, Octopamine HCL, and Ivermectin were purchased from Sigma-Aldrich. Imaging CellCarrier plates were obtained from Perkin Elmer. SYTOX Green, brass sieves, and Breath-Easy^®^ plate seals were from ThermoFisher.

### Nematode culture

Nematodes were cultured on plant hosts growing in germination pouches within temperature-controlled growth chambers set for a 16/8-hour light/dark cycle and ambient humidity at 27 ºC. Soybean Cyst Nematode (SCN) eggs were harvested from soybean plants, cv. Williams 82, 4 weeks after inoculation of Second Stage Juveniles (J2s) on 1-week old seedlings. Eggs were isolated from cysts using the standard graduated sieve system starting at 850µm to final egg capture on a 25µm sieve^10,21^, and captured eggs were placed on two layers of laboratory tissues supported by a wire screen sitting on a Petri dish filled with MilliQ water touching the bottom of the screen. This allowed hatched J2s to migrate through the laboratory tissue and fall into the petri dish. After an overnight incubation, J2s were collected to re-inoculate soybean plants or for use in treatments. Root knot Nematodes (RKN) were cultured in a similar fashion, but on cucumber plants. For egg collection of RKN, root galls were minced and shaken for 3 minutes in a 0.25% sodium hypochlorite solution and run through a series of sieves until final collection on a 25µm sieve. This assured complete rinsing and removal of residual sodium hypochlorite^22^. The RKN J2s were then collected and used in the same fashion as described for the SCN.

### Worm bulk staining

For each batch staining of worms, approximately 100,000 J2 stage nematodes were used for both species. J2s were collected in DI water and concentrated via gravity settling, and subsequently diluted to obtain no more than 15,000 J2/mL. 1 mL aliquots of the resulting worm suspensions were centrifuged at 2,000 rpm for 5 minutes, followed by removal of the resulting supernatant. PKH26 dye was prepared following the manufacture’s recommendations to a concentration of 30µL of dye per mL of buffer, and 1mL of the dye solution was added to each tube of worms. Worms were resuspended by vortexing, followed by a 5-minute incubation in the dark. Following dye incubation, an additional centrifugation step was performed to remove the dye buffer, and worms were washed 3 times via centrifugation in MilliQ water containing 1% bovine serum albumin.

### Microbial supernatant preparation

All microbial broth sample preparation was performed in 24 or 96 well plate formats. A diverse set of ∼2500 novel soil microbial isolates were grown for 3-7 days at 25 ºC. After growth, samples were centrifuged, followed by filtration of the supernatant through a 0.45µm filter. Samples were then stored at -80ºC and assayed as described.

### Assay Set-up

After PKH26 worm staining was completed, stained worms were counted and diluted with MilliQ water and 0.01% Tween to a concentration of 1 worm/µL to prepare them for addition to the assay plates. Stock formulated antimicrobial Penicillin/Streptomycin/Amphotericin B and SYTOX Green dye were added to the worm stock at a final concentration of 20,000 units/mL penicillin, 20mg/mL streptomycin sulfate, 50µg/mL amphotericin B, and 10µM SYTOX Green. Using a Multidrop (Thermofisher), 50µL of the J2 solution were added to each well of a 96-well, black, clear-bottom view plate, slowly agitating the stock during dispensing to keep distribution of J2s regular. 50µL of test sample was then added to each well using liquid handling robotics. Samples were treated in triplicate, and each assay plate contained 8 negative control wells, 4 wells of 0.1% NaOH as the positive control for mortality, and 4 wells of 10mM Ivermectin as the positive control for non-movement. Breathable plate seal membranes (Breath-Easy^®^) were added to each plate to reduce liquid evaporation during incubation. Plates were then incubated for 48 hours at 27^0^C. After incubation, the Breath-Easy membranes were removed from the plate, and 10µL/well of a 10mM octopamine solution in MilliQ water was added to each plate. Plates were then imaged within 24 hours on an INCell 2200. Image acquisition was done using a 2X objective, and sequential Cy3, FITC, Cy3 exposures. A 3 second delay was added between the FITC and final Cy3 exposure.

### Algorithm development and data reporting

Object detection was developed using the Cy3 signal from the worm cuticle staining with PKH26. To eliminate noise, area, length, and intensity thresholding was used, with values unique to each species. Motility was determined by overlaying the worm signal at time 0 and time 3 sec. A worm that completely moved out of location is represented with a movement signal of 0, while a completely unresponsive, non-moving worm value is 1. Each individual worm movement score was assigned a classification of “still” or “move”, and the total scored worms for each treatment well were averaged to assign a classification for that particular treatment. “Still” was defined as a movement score of greater than 0.869 for RKN and greater than 0.9 for SCN. SYTOX Green signal area was detected in the FITC channel, using size and intensity gating. Positive mortality was measured if a significant area of the worm, as defined by the PKH26, was colocalized with the SYTOX Green signal area. All algorithm settings are available upon request.

After analysis, analyzed image data was output as well summary data for total worm counts for both time points, total count of dead worms, and total intensity levels for both FITC and Cy3 channels. Percent effect for each sample was calculated based on the percent of worms per well affected by the microbe sample using a Python script for data analysis (Supplemental Table 2). Final microbial effect was reported as an average of the well replicates based on the following formulas:

Raw (unadjusted) motility and mortality fractions were first calculated for each well as: 
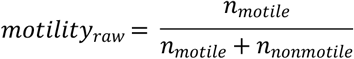

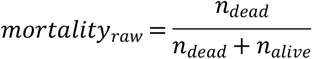
where n represented nematodes in the well. These raw fractions were then used to calculate the control-adjusted motility and mortality for each microbial sample: 
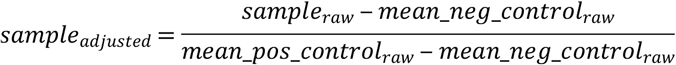

Controls specific to each analysis and subsequent calculations are shown in the table and equations below. Note that motility expresses proportion of nematodes still motile, while mortality expresses proportion of nematodes dead.

|  | Motility | Mortality |
| --- | --- | --- |
| Negative control | NaOH (dead = immobilized) | Untreated (live) |
| Positive control | Untreated (potentially motile) | NaOH (dead) |
| Industry standard | Ivermectin (likely immobilized) |  |

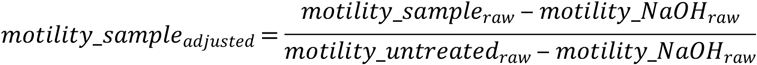

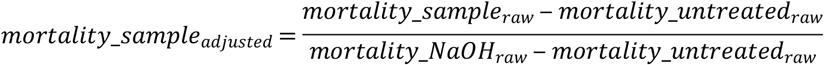

Adjusted motility and mortality scores were converted to percentages and averaged across replicate plates to provide final percent effect scores for each microbe sample.

## Results and Discussion

The assay we developed utilized two different fluorescent dyes, the lipophilic membrane PKH26 dye^9^ which was used to pre-stain large pools of worms collected from nematode cultures maintained on plant hosts grown in germination pouches^10^, and the viability dye SYTOX Green^11^ which had been previously shown to stain non-viable nematodes^12^. By fluorescently pre-staining the nematodes with the permanent fluorescent marker PKH26, we could produce high contrast images that allowed identification and localization of individual worms in treatment wells. Linking influx of SYTOX Green to each worm then allowed us to determine the viability of each individual in the treated population. Additionally, by utilizing the time-lapse capabilities of our HCA instrument along with image analysis, we could measure the movement of each worm down to micron levels. All the imaging was performed on the GE INCell 2200 instrument platform, a high content platform widely used in the human drug discovery area that allows cellular phenotypic analysis, individual object detection and segmentation, as well as movement analysis^13–15^.

Assay validation experiments focused on several variables. First, the stability of PKH26 dye staining of the worms over multiple days in treatment plates and multiple imaging exposures was examined to assure minimal toxicity of the dye itself on the worms as well as minimal photobleaching of PKH26 when several exposures were performed. Secondly, worm distribution into 96-well test plates was optimized to assure accurate software individual object detection and phenotyping (Figure 1). Finally, since the ultimate goal of our screening paradigm was to identify microbe strains that exuded molecules capable of modulating the behavior of or killing the infective Second Stage Juvenile (J2) of RKN and SCN nematodes, we wanted to include a control in each treatment plate as an indicator of mortality so that we could easily distinguish between living, paralyzed, and dead worms. By taking advantage of the epi-fluorescent capabilities of the INCell 2200, we were able to add a viable dye as part of our assay readout. Our final protocol utilized SYTOX Green penetration in worms treated with 0.1% NaOH for this purpose (Figures 1B and 1C). We also included the well-characterized anthelmintic compound Ivermectin^16^ as a control for identification of paralyzed worms (Figure 1C). To demonstrate the reproducibility and accuracy of the selected controls, Z’-factors^17^ were calculated for the differences between both positive control treated worms and negative control worms (Figure 2, Table 1).

**Figure 1.**
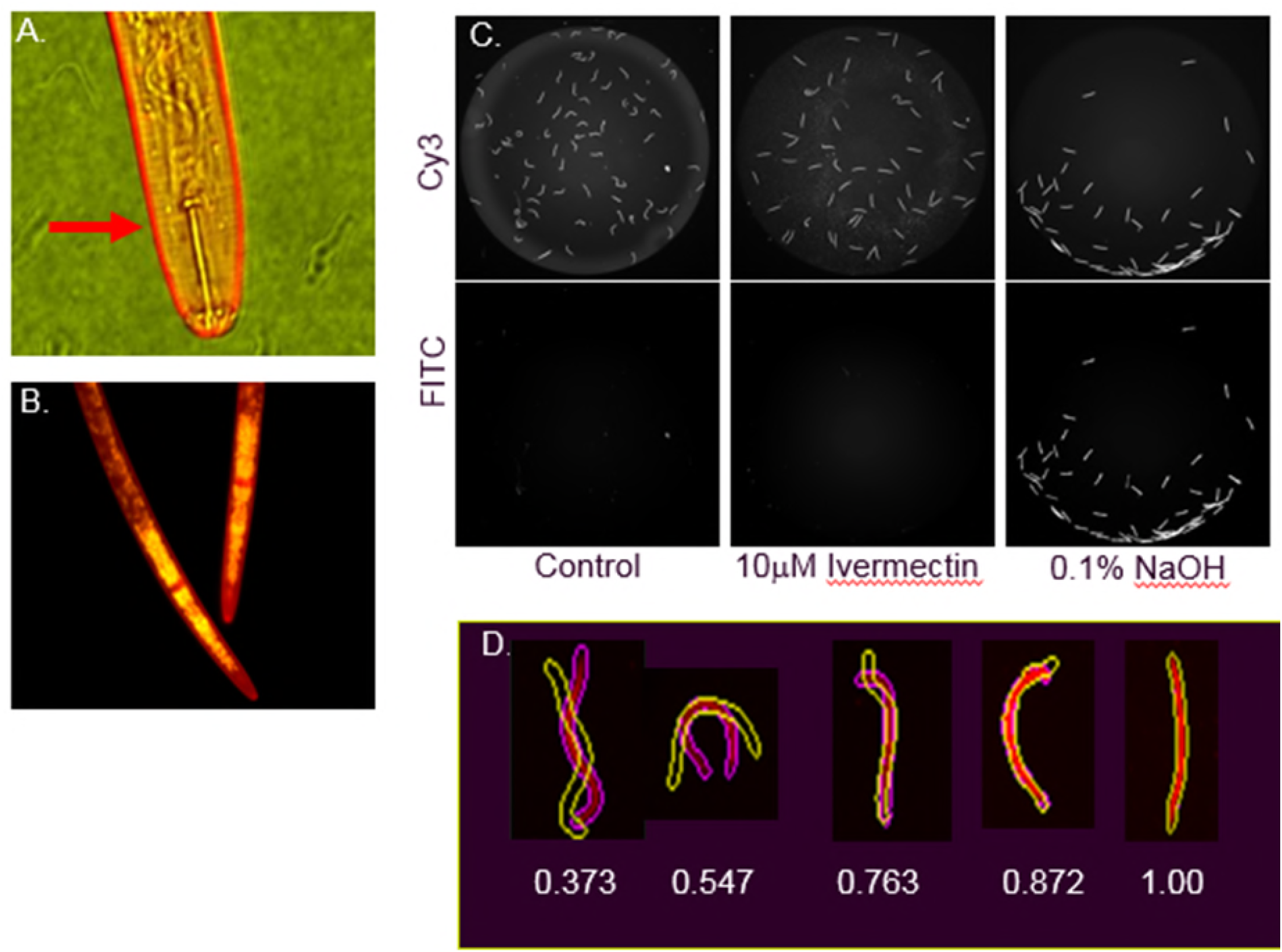
Imaging and quantification of nematode viability and movement. All images were obtained with a GE 2200 HCA instrument. (**A**.) PKH26 lipid dye stains the outer cuticle of SCN, indicated by red arrow. (**B**.) A representative 60X merged image of heat-shocked RKN stained with both PKH26 (red) and SYTOX Green (green), showing internal DNA-binding of SYTOX Green after worm death. (**C**.) Representative RKN grey-scale images of PKH26 (Cy3) and SYTOX Green (FITC) treated with 10µM Ivermectin and 0.1% NaOH. Only worms treated with 0.1% NaOH stain with the viable dye SYTOX Green. All intensity values were normalized between sample wells. (**D**.) Examples of Cy3 detected object outlines separated by a 3 second delay and overlaid by the GE Developer analysis software (time 1=solid purple outline, time 2=yellow outline). Movement is ranked based on 100% overlap (given a value of 1) and no overlap (given a value of 0). Fine differences are indicated by assigned score by the analysis software.

**Figure 2.**
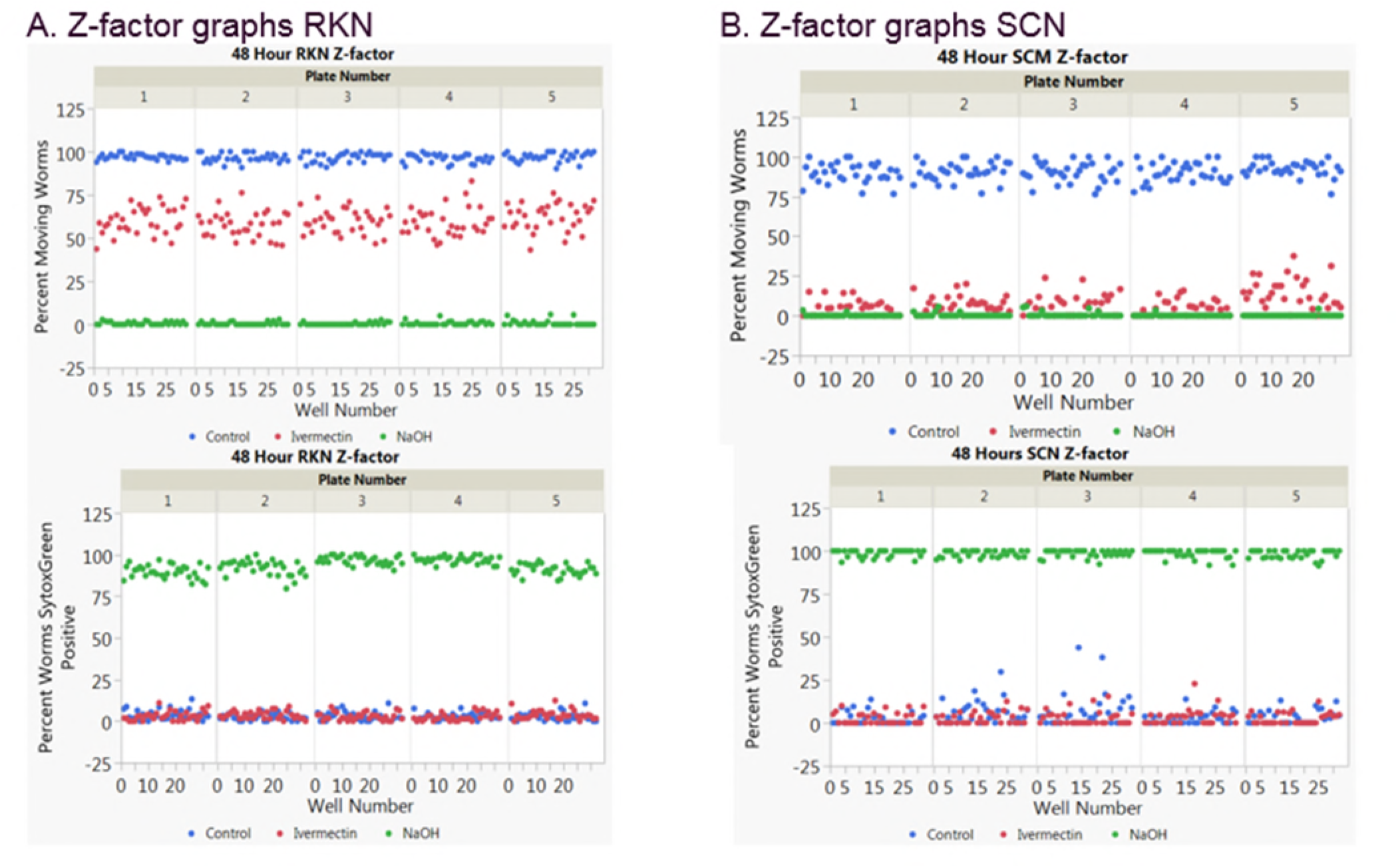
Representative data from a single day of a 3-day set of experiments. For each day, 5 individual 96-well plates were treated with control, 10µM Ivermectin, and 0.1% NaOH, 32 individual wells per treatment plate. For statistical analysis, Z’-factors were calculated for each individual plate using the following formula: Z’=1-(3*SD Treatment 1 mean + 3*SD Treatment 2 mean)/(Treatment 1 mean – Treatment 2 mean) with control wells being used in all calculations for one of the treatment sets. Each dot represents the data from an individual treatment well. The data for each treatment and plate group well in the outputted visualization, indicating assay robustness.

**Table 1:** Calculated average Z-factors of the 3 days of 5 plates (n=15 plates total representing 480 wells) for mortality based on 0.1%NaOH (**A**.), and movement for both 10µM Ivermectin (**B**.) and 0.1%NaOH (**C**.). Images were captured at 24 hours, 48 hours and 6 days of treatment and all images transformed with Cell Developer software based on the algorithm described in the **Methods** section. Percent CV was calculated from the mean calculated Z’-factor for each individual day. To maximize worm exposure to treatment, and minimize variation introduced over treatment time, all final assays were performed using the 48-hour time point for data acquisition.

|  |  |  |  |
| --- | --- | --- | --- |
| **A.**<br>**Calculated percent mortality from 0.1% NaOH day-to-day Z'-factor** | **24 hour RKN** | **48 hour RKN** | **6 day RKN** |
| Ave. percent mortality | 0.88 | 0.85 | 0.62 |
| Standard deviation | 0.04 | 0.07 | 0.33 |
| Percent CV | 4.90 | 8.04 | 52.83 |
|  | **24 hour SCN** | **48 hour SCN** | **6 day SCN** |
| Ave. percent mortality | 0.85 | 0.76 | 0.41 |
| Standard deviation | 0.06 | 0.04 | 0.16 |
| Percent CV | 6.82 | 5.20 | 38.07 |
| **B.**<br>**Calculated percent movement inhibition from 10µM Ivermectin day-to-day Z'-factor** | **24 hour RKN** | **48 hour RKN** | **6 day RKN** |
| Ave. percent movement | 0.67 | 0.05 | -2.71 |
| Standard deviation | 0.07 | 0.21 | 4.04 |
| Percent CV | 11.17 | 392.55 | -149.16 |
|  | **24 hour SCN** | **48 hour SCN** | **6 day SCN** |
| Ave. percent movement | 0.45 | 0.59 | 0.31 |
| Standard deviation | 0.14 | 0.05 | 0.22 |
| Percent CV | 31.66 | 8.20 | 69.78 |
| **C.**<br>**Calculated percent movement inhibition from 0.1% NaOH day-to-day Z'-factor** | **24 hour RKN** | **48 hour RKN** | **6 day RKN** |
| Ave. percent movement | 0.91 | 0.89 | 0.71 |
| Standard deviation | 0.01 | 0.01 | 0.20 |
| Percent CV | 0.79 | 1.31 | 28.88 |
|  | **24 hour SCN** | **48 hour SCN** | **6 day SCN** |
| Ave. percent movement | 0.75 | 0.74 | 0.79 |
| Standard deviation | 0.04 | 0.09 | 0.04 |
| Percent CV | 5.76 | 12.58 | 4.84 |

Our final assay conditions and detection algorithms were able to distinguish test samples capable of only paralyzing worms from samples capable of killing worms (Figure 1C). To assure we measured movement in all worms capable of movement responses, immediately before image capture the neurotransmitter Octopamine was added to all sample wells^18^. This caused a generalized increase of movement in the worm population. We then developed a detection algorithm within the GE Cell Developer software that allowed for the determination of worm movement using a timed approach, capturing two images separated by 3 seconds. The advantage of our HCA system and protocol was the coupling of high resolution timed image capture with plate movement control and multiple fluorescent readouts resulting in a high throughput phenotypic screening method with separate measurements for worm movement and worm mortality. Each individual well of our 96 well assay plates had 3 separate images captured in the following sequence, Cy3 (PKH26 dye), FITC (SYTOX Green dye), and a second Cy3 image captured after a 3 second delay. The two Cy3 images were then overlaid using the colocalization function via the Cell Developer analysis software. Total worm movement that took place between the two-timed images could be quantified at single pixel resolution. In our case, at 2X magnification, that resolution is 3.25µm. Our final movement readout was based on the percentage of total overlapping pixels for individual worms (Figure 1D), with each worm assigned a movement score by the analysis software. Movement scores were then averaged for total worms in each treated well, followed by classification of each treatment well as “movement” or “still”.

Using our colocalization measurements on individual worms, we identified a dose-dependent range of movement for Ivermectin-treated RKN and SCN. We used this scale to calibrate our paralysis-acceptability limits, as slight movement could still be seen in Ivermectin-treated worms. Calculated EC_50_ movement values changed over time and were different for each species (Table 2 and Figure 1B). We did not observe any significant mortality associated with continuous 48-hour Ivermectin treatment for any dose. Concentrations of Ivermectin greater than 10µM resulted in fluorescent precipitation in the assay wells, reducing our ability to determine mortality effects with SYTOX Green above 10µM Ivermectin. As a comparison, Ferreira et. al. reported the IC_50_ of Ivermectin treated *C. elegans* for mortality as 261.4µM, but the EC_50_ for movement was 0.87µM, a ∼300-fold difference in measured endpoints^12^.

**Table 2:** Calculated EC_50_ values for Ivermectin and NaOH. All treatments were performed in 3 individual 96 well plates, n=8 wells for each individual concentration, for 48 hours. EC_50_ values were calculated with GraphPad Prism (version 7.04) software using Nonlinear regression curve fit. As expected, we were unable to calculate an EC_50_ for Ivermectin-driven mortality.

| **Percent Mortality** | **RKN 48 hours** | **SCN 48 hours** |
| --- | --- | --- |
| µM Ivermectin EC<sub>50</sub> | **>40µM** | **>40µM** |
| µM Ivermectin EC<sub>50</sub> %CV | **NA** | **NA** |
| % NaOH EC<sub>50</sub> | **0.01%** | **0.006%** |
| % NaOH EC<sub>50</sub> R square | **0.995** | **0.972** |
| **Percent Movement** |  |  |
| µM Ivermectin EC<sub>50</sub> | **0.43µM** | **1.06µM** |
| µM Ivermectin EC<sub>50</sub> R square | **0.801** | **0.980** |
| % NaOH EC<sub>50</sub> | **0.03%** | **0.02%** |
| % NaOH EC<sub>50</sub> R square | **0.994** | **0.758** |

Because the images are a permanent record of what is in each well at the time of image acquisition, we were able to compare manually determined worm phenotypes from the images to machine-determined phenotypes and found a very high level of agreement (Table 3). Based upon the robust statistically significant Z-factor^14, 15^ and reproducible EC_50_ results obtained for control treatments during our method development, we have concluded that the automated image analysis protocol for nematode high throughput screening described here is robust, reproducible and generates dependable data without variation inherent in manual worm observation (Figure 2 and 3, Tables 1 and 2). Inherent with the automated focus, plate movement and stage movement of HCA instrumentation is the ability to screen large sample libraries in a short time. We typically screened up to 60 individual 96-well plates at a time, each plate taking approximately 10 minutes to capture all images, for a total run time of 10 hours for 5760 individual analyzed wells. This level of sample testing throughput should allow for faster discovery and development of urgently needed anti-nematode chemistries or microbial inoculants that will help achieve the goal of reducing nematode-related crop yield loss and increasing food security. Additionally, it seems reasonable to assume that similar approaches as those presented here should be applicable to both relevant animal and human nematode and related parasites. For example, with growing resistance becoming apparent in veterinary applications of Ivermectin^19,20^, additional animal health treatment options has become a necessity.

**Figure 3.**
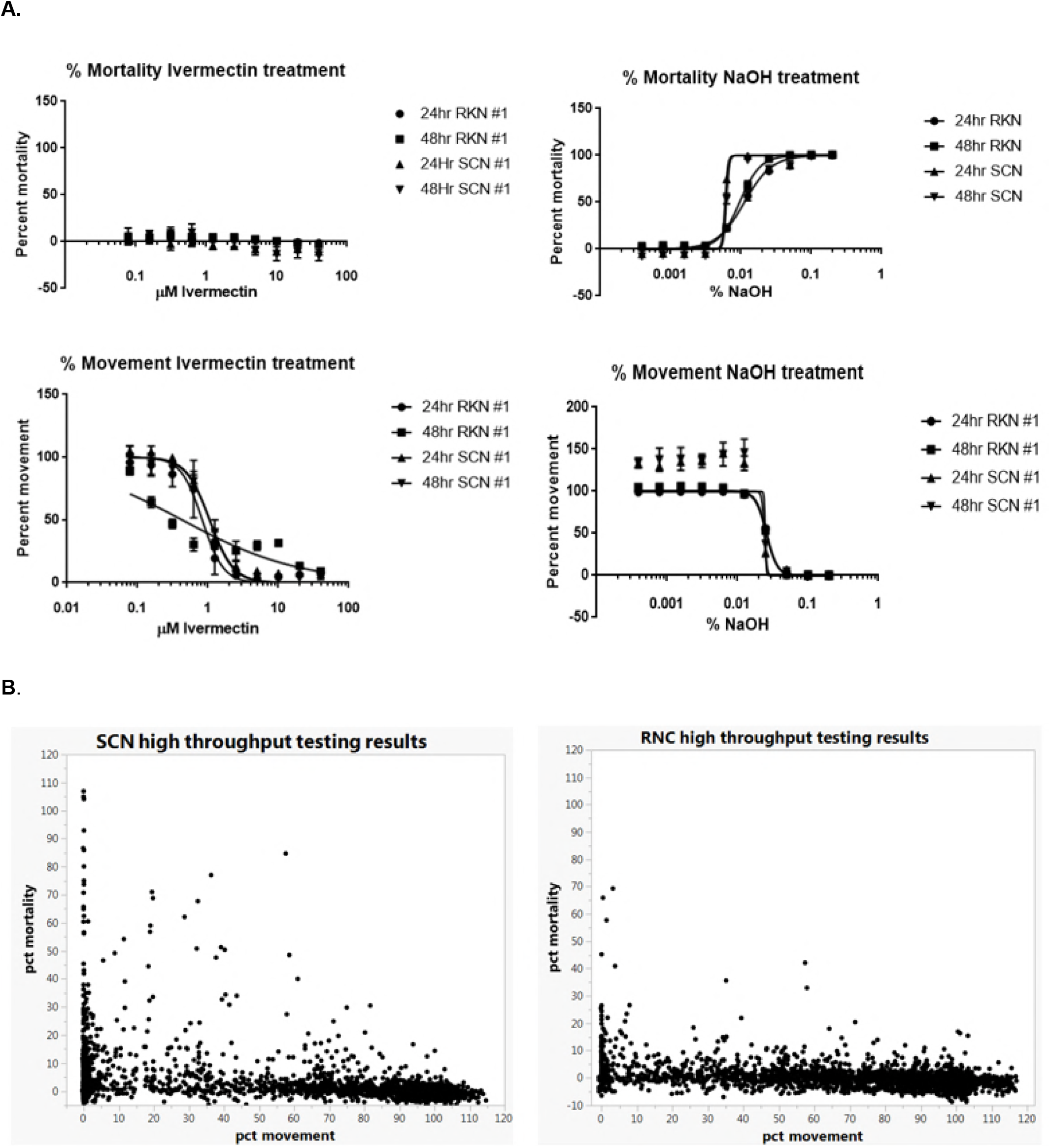
Assay accuracy and reproducibility. **(A.)** Demonstration of accuracy and repeatability based on 3 separate plates containing 8 individual EC_50_ determinations for Ivermectin and NaOH. Each concentration displayed is the mean of the individual dose responses on each plate, with error bars indicating standard deviation between the replicate plates. We observed no effect from Ivermectin on mortality for either species but noted reproducible EC_50_ values for the percent movement measurement in both species. For NaOH treated nematodes, both percent movement and percent mortality measures were calculated for both species (see Table 1 for mean calculated EC_50_ values). **(B.)** Results from the high throughput microbial supernate screening assays. Approximately 2,500 individual isolates were grown for 3-7 days, and the spent media tested for the ability to influence nematode movement or mortality. Our results indicate a bias in our microbial library for strains that effect movement at a level well above strains that influence worm mortality. Additionally, as expected, as percent movement drops, the number of strains showing a higher percent mortality measurement increases.

**Table 3:** Manual counting vs. machine counting results. Images of nematodes were manually scored and compared to scored achieved using the developed automated scoring process. To achieve mixed viable/non-viable samples, a pool of worms were heat-killed via treatment at 90°C for 10 minutes. This heat-killed pool was then mixed with a pool of control worms. Following the described methods, images were captured for 6 different sample wells containing ∼50-70 total worms per well.

| Treatment sample | Percent mortality detected automated algorithm | Percent mortality detected human counting |
| --- | --- | --- |
| Well #1 | 45 | 42 |
| Well #2 | 49 | 61 |
| Well #3 | 27 | 29 |
| Well #4 | 54 | 57 |
| Well #5 | 28 | 22 |
| Well #6 | 57 | 57 |

## Conclusion

The findings presented here indicate that time-lapse capabilities of image analysis are useful for analysis of various biological samples. While nematodes were used here to detect samples that inhibited movement, loss of movement of other organisms, or even single cells, in response to various stimuli could be detected. Instead of loss of movement, stimulation of movement could also be detected. Our studies have also demonstrated that additional parameters (here, viability staining of the nematodes) can be used along with the time-lapse movement detection. There also is the capacity to add additional metrics to truly multiplex the assay platform as described and help elucidate additional biological complexity beyond the described endpoints, further efforts will follow this course.

## Acknowledgements

The authors would like to thank the Novozymes BioAg Discovery team for access to the test microbe strains, and we would like to especially thank Dr. Louisa Liberman for her critical reviews of our draft manuscripts.

